# Bile acid composition regulates the manganese transporter Slc30a10 in intestine

**DOI:** 10.1101/2020.02.11.944124

**Authors:** Tiara R. Ahmad, Sei Higuchi, Enrico Bertaggia, Allison Hung, Niroshan Shanmugarajah, Nicole C. Guilz, Jennifer R. Gamarra, Rebecca A. Haeusler

## Abstract

Bile acids (BAs) comprise heterogenous amphipathic cholesterol-derived molecules that carry out physicochemical and signaling functions. A major site of BA action is the terminal ileum, where enterocytes actively reuptake BAs and express high levels of BA-sensitive nuclear receptors. BA pool size and composition are affected by changes in metabolic health, and vice versa. One of several factors that differentiate BAs is the presence of a hydroxyl group on C12 of the steroid ring. 12a-hydroxylated BAs (12HBAs) are altered in multiple disease settings, but the consequences of 12HBA abundance are incompletely understood. We employed mouse primary ileum organoids to investigate the transcriptional effects of varying 12HBA abundance in BA pools. We identified *Slc30a10* as one of the top genes differentially induced by BA pools with varying 12HBA abundance. SLC30A10 is a manganese (Mn) efflux transporter critical for whole-body manganese excretion. We found that BA pools, especially those low in 12HBAs, induce cellular manganese efflux, and that *Slc30a10* induction by BA pools is driven primarily by lithocholic acid signaling via the vitamin D receptor. Administration of lithocholic acid or a vitamin D receptor agonist resulted in increased *Slc30a10* expression in mouse ileum epithelia. These data demonstrate a previously unknown role for BAs in intestinal control of Mn homeostasis.

---

Bile acids (BAs) are cholesterol catabolites that regulate many biological functions, including multiple aspects of macronutrient metabolism. One of the mechanisms by which they do so is by promoting lipid emulsification and absorption (1–3). A second mechanism is by acting as a ligand for BA receptors, which can regulate lipid and glucose metabolism (1–4). It is underappreciated that there is structural diversity among BAs that results in variable capacities to activate BA receptors (5–10). This structural diversity arises from the number and position of hydroxyl groups and the conjugation of the molecule to glycine, taurine, or neither (11, 12). Thus the composition of the BA pool may affect the activity of BA receptors. However, the biological consequences of altered BA pool composition are not fully known.

One key determinant of BA composition is the hepatic enzyme sterol 12α-hydroxylase (encoded by *CYP8B1*). By adding a 12α-hydroxylation to an intermediate of the BA synthesis pathway, CYP8B1 determines the hepatic synthesis of cholic acid (CA) instead of chenodeoxycholic acid (CDCA) (Fig. 1A) (12, 13). In settings of insulin resistance, there is an increased proportion of CA, its bacterial metabolite deoxycholic acid (DCA), and their conjugates—collectively termed 12α-hydroxylated BAs (12HBAs) in the BA pool (14, 15). Rates of 12HBA synthesis and CA conversion to DCA are also higher in insulin resistance and type-2 diabetes (16, 17). Moreover, even in healthy subjects, increases in 12HBAs are correlated with the characteristic metabolic abnormalities of insulin resistance (14). In contrast, *Cyp8b1*^*–/–*^ mice, which lack 12HBAs, are protected from western-type diet-induced weight gain and atherosclerosis compared to wild-type mice (18–20). *Cyp8b1*^*–/–*^ mice also show improved glucose tolerance, which has been proposed to be due to increased secretion of glucagon-like peptide-1 (GLP-1) (21). Mice subjected to vertical sleeve gastrectomy (VSG), a common weight loss surgery, exhibit reduced *Cyp8b1* mRNA expression as well as lower ratios of 12HBAs:non-12HBAs in circulation (22). Furthermore, siRNA against *Cyp8b1* improved non-alcoholic steatohepatitis in mice (23). Thus, CYP8B1 inhibition is a potential therapeutic target for metabolic diseases. However, the biological processes that are regulated by 12HBAs are incompletely understood.

**Figure 1.**
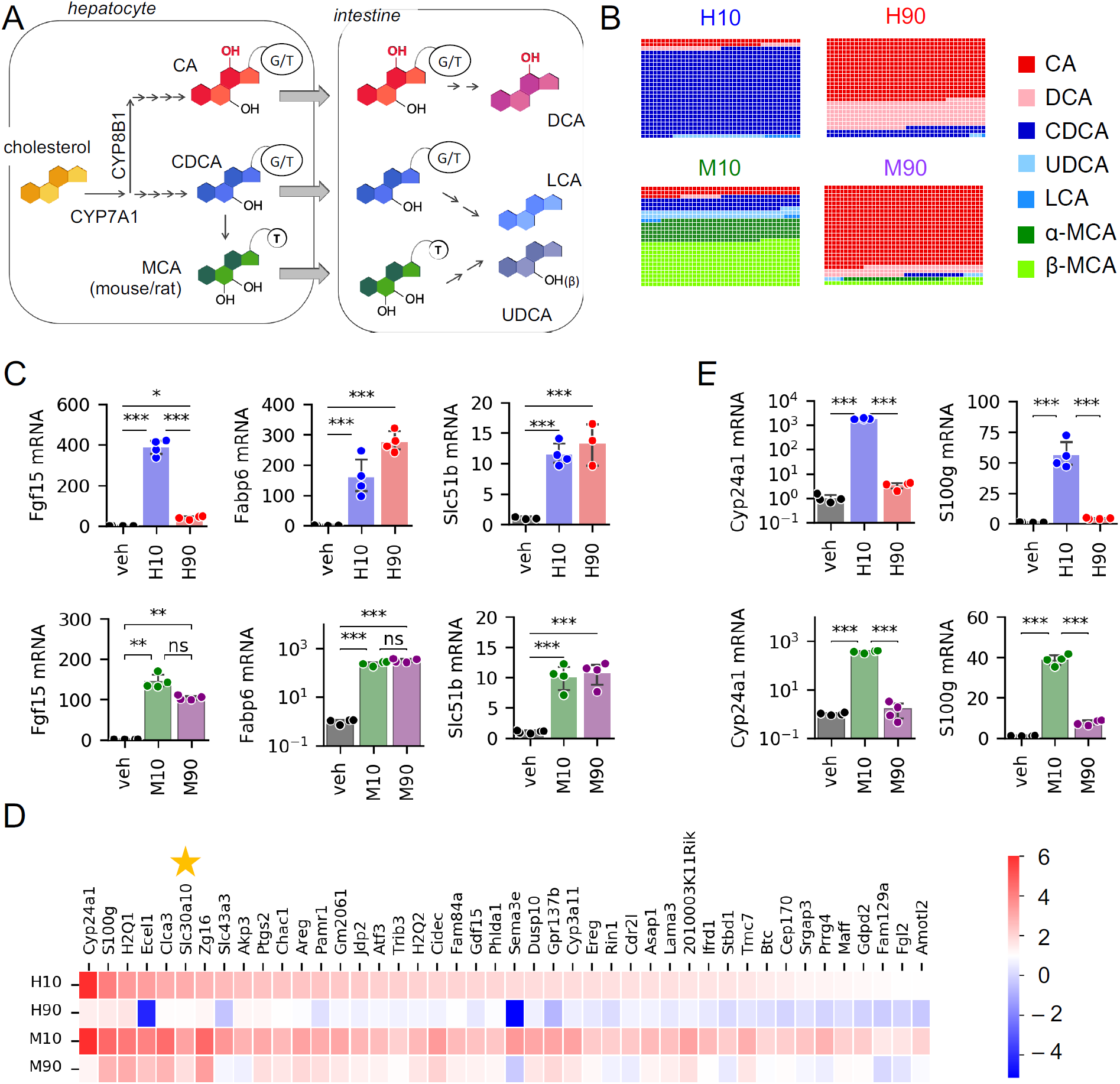
Differential regulation of gene expression by BA pool composition. (A) Simplified BA synthesis pathway. (B) Composition of BA pools used in this study. Glycine-conjugated BAs were used in human pools, while taurine-conjugated BAs were used in mouse pools, except LCA, which was unconjugated. (C) Expression of FXR target genes measured by qPCR. n=3-4 wells of organoids per condition. (D) Genes preferentially induced by H10 and M10 compared to H90 and M90 identified by RNAseq (log2 fold-change > 1 and *p*_adj_ < 0.05). n=3 per condition. (E) Expression of VDR target genes measured by qPCR. n=3-4 wells per condition. \**p*<0.05, \*\**p*<0.01, and \*\*\**p*<0.001.

A major site of BA signaling is the intestine, which encounters high BA concentrations, approximately 2-12 mM after a meal (24, 25). The intestine epithelium expresses at least three BA receptors. The transcription factor FXR regulates BA transport and feedback suppression of hepatic BA synthesis, and also modulates lipid and glucose metabolism (1). The membrane receptor TGR5 regulates glucose homeostasis and colonic motility by promoting the secretion of GLP-1 and serotonin (26–28). Another intestinal receptor responsive to certain BAs is the vitamin D receptor (VDR), whose canonical role is to promote calcium and phosphate absorption (29). For each of these receptors, the best reported endogenous BA agonists are non-12HBAs. For FXR it is CDCA (5, 6, 9), for TGR5 and VDR it is lithocholic acid (LCA) (7, 8, 10). LCA is formed by 7α-dehydroxylation of CDCA by bacterial enzymes in the gut. Thus BA composition is predicted to impact signaling through multiple receptors in the intestine.

Investigating the effects of BA composition on intestinal BA signaling *in vivo* is challenging because of the continued presence of endogenous BAs. This is particularly important for experiments in mice, as mice contain a class of BAs–muricholic acids (MCAs), which are non-12HBAs–that are not found in healthy adult humans. To fill the gap, we used primary murine intestinal organoids, also called enteroids. These organoids are generated from stem cells of the intestinal crypts, and contain all known cell types of the intestinal epithelium (30). We investigated the effects of BA pools of different compositions on ileal organoids, with a particular focus on the effects of lowering 12HBAs (to mimic CYP8B1 inhibition). Furthermore, we addressed the interspecies differences in BAs by using BA pools that we designed to mimic the effects of CYP8B1 inhibition in humans and mice.

We unexpectedly found that varying 12HBA proportions modulates expression of *Slc30a10*, a manganese (Mn) efflux transporter critical for whole-body Mn excretion. Cellular Mn levels are tightly regulated, as Mn is essential for numerous cellular processes, yet its excess is toxic (31). Our data demonstrate a previously unknown role of BAs in intestinal control of metal homeostasis.

## Results

### Distinct BA pool compositions differentially induce gene expression

To test the effects of a low 12HBA pool on intestinal gene expression, we designed four distinct BA pools with which to stimulate primary murine ileal organoids. The differences between the pools were due to two key features: (i) the proportion of 12HBAs, either 10% (low) or 90% (high), and (ii) the BA pool of the species we model, either human or mouse (Table 1, Fig. 1B). In the human pools, there was a larger proportion of DCA and the BAs were glycine-conjugated, whereas in the mouse pools, BAs were taurine-conjugated, to mimic the natural abundance in those species (4, 32, 33). Mouse BA pools contained MCAs, whereas these were not included in the human pools. Thus, the four BA pools are labeled human low 12HBA (H10), human high 12HBA (H90), mouse low 12HBA (M10), and mouse high 12HBA (M90). Importantly, the total BA pool concentration was the same across all treatment groups. We prepared the four BA pools in mixed micelles containing oleic acid, 2-palmitoyl glycerol, phosphatidylcholine, and free cholesterol, to mimic conditions of the intestinal lumen. We used the four lipid-emulsified BA pools to treat primary murine ileal organoids. The vehicle control contained all micelle components except BAs. We performed bulk RNA sequencing after 24 hours of treatment.

**Table 1.** Composition of BA pools.

| BA group | BA species | mol % in H10 | mol % in H90 | mol % in M10 | mol % in M90 |
| --- | --- | --- | --- | --- | --- |
| 12HBA | Cholic acid (CA) | 7 | 63 | 9.0 | 81.0 |
| 12HBA | Deoxycholic acid (DCA) | 3 | 27 | 1.0 | 9.0 |
| Non-12HBA | Chenodeoxycholic acid (CDCA) | 86.85 | 9.65 | 13.5 | 1.5 |
| Non-12HBA | Ursodeoxycholic acid (UDCA) | 2.25 | 0.25 | 8.1 | 0.9 |
| Non-12HBA | Lithocholic acid (LCA) | 0.90 | 0.10 | 0.9 | 0.1 |
| Non-12HBA | α-Muricholic acid (α-MCA) | | | 22.5 | 2.5 |
| Non-12HBA | β-Muricholic acid (β-MCA) | | | 45.0 | 5.0 |

BAs were in their glycine-conjugated forms for human BA pools and taurine-conjugated forms for mouse BA pools, except LCA, which was unconjugated.

We focused on the effects of the low 12HBA pools, as this would mimic the effects of CYP8B1 inhibition. We examined all genes that were significantly induced compared to vehicle, setting thresholds of log_2_FC>1.0 and *p*_adj_<0.05 for differential expression. The low 12HBA pools collectively induced 1361 genes. Of these, 516 reached those thresholds for both H10 and M10 pools, 95 reached those thresholds for H10 only, and 750 for M10 only (Supp. Fig. 1A). Pathway analysis indicates enrichment of genes involved in lipid metabolism, consistent with known effects of BAs (Table 2).

**Table 2.**
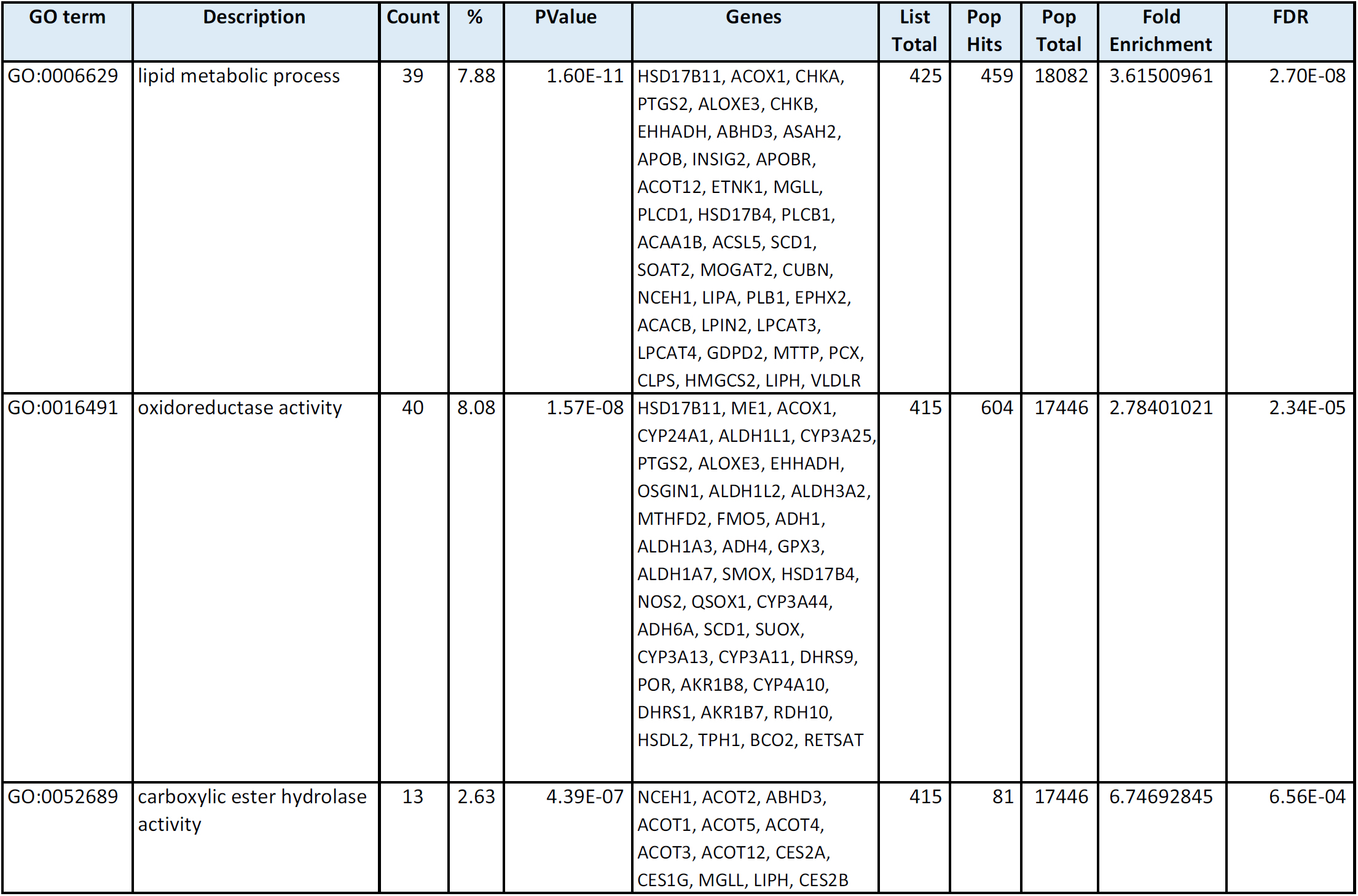
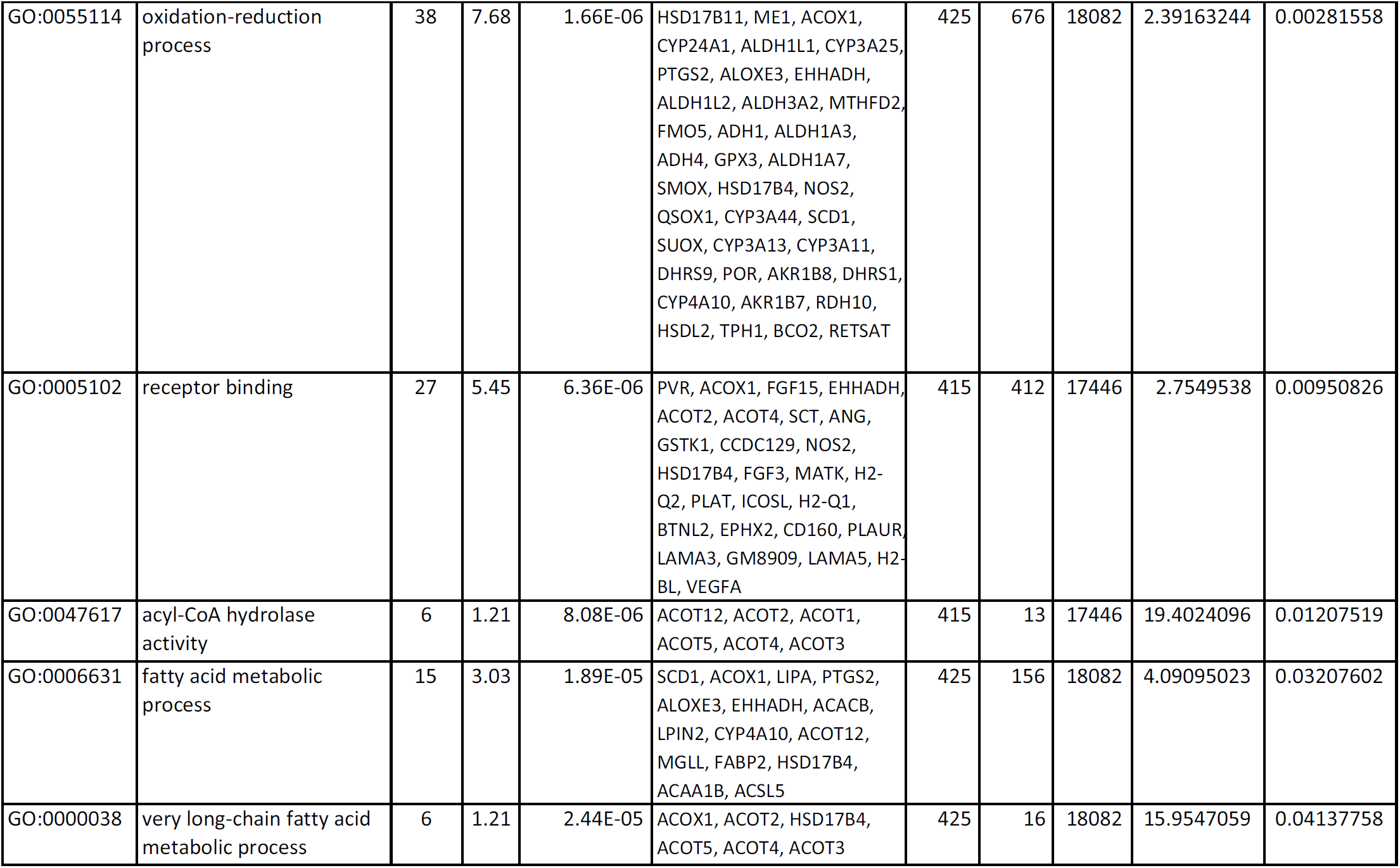
Pathway analysis of 516 genes induced by both human low 12HBA and mouse low 12HBA pools.

Next, we focused on the subset of these genes that are differentially regulated by low-versus high-12HBA pools, with a particular focus on those for which the effects were shared between human and mouse pools. We used a threshold of log_2_FC>1.0 and P_adj_<0.05 for differential expression. The majority of genes did not reach this threshold, indicating that they are similarly regulated by both low- and high-12HBA pools, or are differentially regulated by low-versus high-12HBAs in human pools only or in mouse pools only (Supp. Fig. 1B). These genes included canonical FXR targets such as *Fgf15, Fabp6* (encoding the ileal bile acid binding protein, Ibabp), and *Slc51b* (encoding the basolateral BA efflux transporter Ostβ), and we validated these by qPCR (Fig. 1C).

There were 44 genes that were preferentially induced by H10 and M10 compared to H90 and M90, respectively (Fig. 1D). Among these, we noted that several are known transcriptional targets of VDR. These included *Cyp24a1, S100g*, and *Cyp3a11*, and we validated these by qPCR (Fig. 1E). This is consistent with the concepts that (i) certain BAs, especially the non-12HBA lithocholic acid (LCA) and its conjugates, can activate VDR in the µM range (10), and (ii) the low 12HBA pools (*i.e.* H10 and M10) contain more LCA (9 µM, as opposed to 1 µM in the high 12HBA pools).

Next, we validated the RNA-seq findings in multiple experimental systems. Using organoids derived from multiple mice, we confirmed that all BA pools induced expression of FXR targets *Fgf15* and *Fabp6* (Supp. Fig. 1C). We also confirmed that VDR targets *Cyp24a1* and *S100g* were preferentially induced by low 12HBA pools (Supp. Fig. 1D). We found that delivery in micelles was not required, and that BAs *per se* were sufficient to induce *Fgf15, Fabp6*, and *S100g* (Supp. Fig. 1E). Lastly, to determine whether differential response to BA pools also occur in human cells, we performed experiments in Caco-2 cells, which are enterocyte-like cells derived from a human epithelial colorectal tumor. We observed that FXR and VDR targets were robustly induced by H10, but not H90 (Supp. Fig. 1F, 1G).

### BA composition regulates Slc30a10 transcription and cellular Mn efflux

Among the genes identified to be differentially expressed between BA pools with varying 12HBA abundance, one of the most robust was *Slc30a10* (Fig. 1D). This gene encodes a Mn efflux transporter critical for whole-body Mn excretion (34–41). Humans with mutations in *SLC30A10* develop hypermanganesemia, accompanied by parkinsonism and cirrhosis (34, 35, 42–44).

In gut organoids, low 12HBA pools were superior to high 12HBA pools in inducing *Slc30a10* (Fig. 2A, 2B). This preferential induction was consistent across multiple independent batches of organoids (Fig. 2C, 2D). This differential expression of *Slc30a10* was also observed when BA pools were delivered without micelles (Fig. 2E), indicating a direct effect of BAs. We also validated that the H10 pool induces *SLC30A10* in human Caco-2 cells (Fig. 2F).

**Figure 2.**
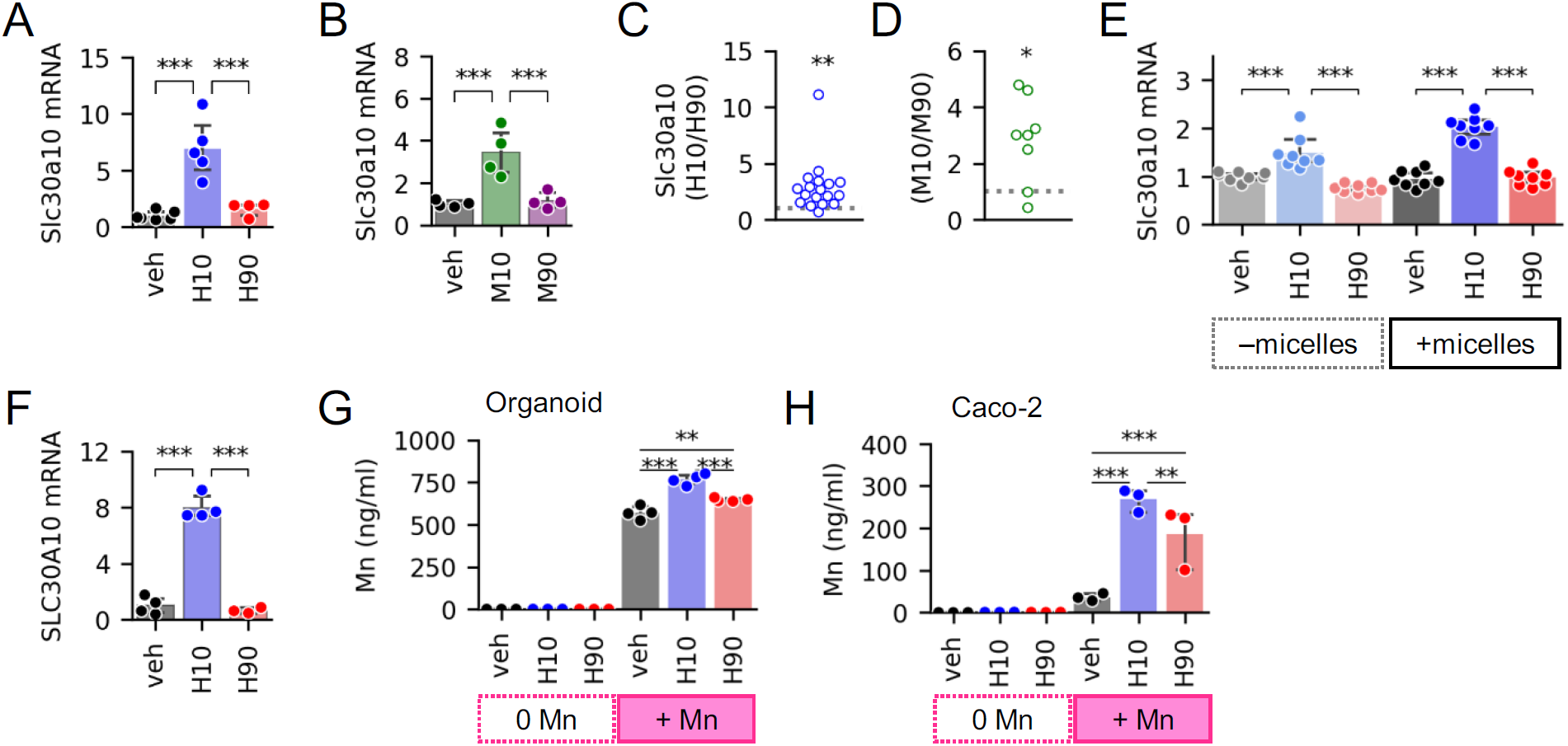
Low 12HBA pools regulate Slc30a10 expression and Mn efflux. *Slc30a10* mRNA levels in gut organoids treated with (A) human BA pools, (B) mouse BA pools. n=4-6 wells of organoids per condition. (C, D) Aggregate fold-change differences in *Slc30a10* induction across different batches of organoids. Each point represents the data derived from a different mouse (n=8-18). The data point plotted is the ratio of (average mRNA levels in low 12HBA-treated organoids [n=3-8 wells])/(average mRNA levels in high 12HBA-treated organoids [n=3-8 wells]). Ratios >1 signify higher expression in low 12HBA-treated group, ratios <1 signify higher expression in high 12HBA-treated group, ratios = 1 signify no difference between low- versus high- 12HBA- treated groups. The gray dotted line marks ratio = 1. \**p*<0.05, \*\**p*<0.01 one-sample t-test with µ=1 (expected ratio for no preferential induction). (E) *Slc30a10* mRNA levels in ileal organoids treated with human BA pools delivered without or with micelles. n=8 wells per condition. (F) *SLC30A10* mRNA levels in Caco-2 treated with human BA pools. n=3-4 wells of cells per condition. (G-H) Mn concentrations in efflux media of (G) organoids and (H) Caco-2 cells, n=3- 4 wells per condition. \**p*<0.05, \*\**p*<0.01, and \*\*\**p*<0.001.

Next we sought to determine whether BA-induced changes in *Slc30a10* transcript levels yield functional cellular consequences. We predicted that the induction of *Slc30a10* expression by low 12HBA pools would increase Mn efflux from organoids. To test this, we carried out Mn efflux assays, where we preloaded cells with Mn, washed away unabsorbed Mn, then treated the cells with vehicle or BA pools, and measured Mn levels in efflux media. Organoids treated with H10 had higher Mn in their efflux media compared to organoids treated with vehicle or H90 (Fig. 2G). Consistently, Caco-2 cells receiving H10 also showed higher Mn efflux compared to vehicle- and H90-treated cells (Fig. 2H). Altogether, these data show that BAs, particularly pools low in 12HBAs, induce cellular Mn efflux, consistent with their induction of *Slc30a10*.

### Slc30a10 transcription is driven by LCA-to-VDR signaling

Recently, *SLC30A10* mRNA expression was found to be induced by the VDR agonist 1α,25-dihydroxyvitamin D_3_ in Caco-2 cells (45). LCA is a potent VDR ligand, with an EC_50_ comparable to CDCA’s EC_50_ for human FXR (10). Although LCA made up less than 1% of the BA pools in our studies, the LCA concentration in the low 12HBA pools was 9 µM. As a point of reference, in duodenum of healthy adult humans, the total bile acid concentration is 20 mM, and LCA makes up 1-3% of the pool, ∼200-600 µM (25, 46, 47). Although LCA is partly sulfated and excreted, a portion is taken up into enterocytes (48, 49). Thus, we investigated whether the differential *Slc30a10* induction by low 12HBA pools was due to LCA-to- VDR signaling.

To test whether LCA is the differentiating factor between H10 and H90 in their induction of *Slc30a10*, we treated ileal organoids with H10 and H90 pools lacking LCA (Supp. Table 1). In pools lacking LCA, we compensated its removal by increasing other non-12HBAs such that the total BA pool concentrations matched. Without LCA, H10 induction of *Slc30a10* was significantly blunted (Fig. 3A). This pattern was similar to that of VDR targets *Cyp24a1* and *S100g* (Fig. 3B). On the other hand, presence or absence of LCA had no effect on FXR targets *Fgf15* and *Fabp6*, consistent with the observation that LCA is not a potent FXR agonist (Fig. 3C).

**Figure 3.**
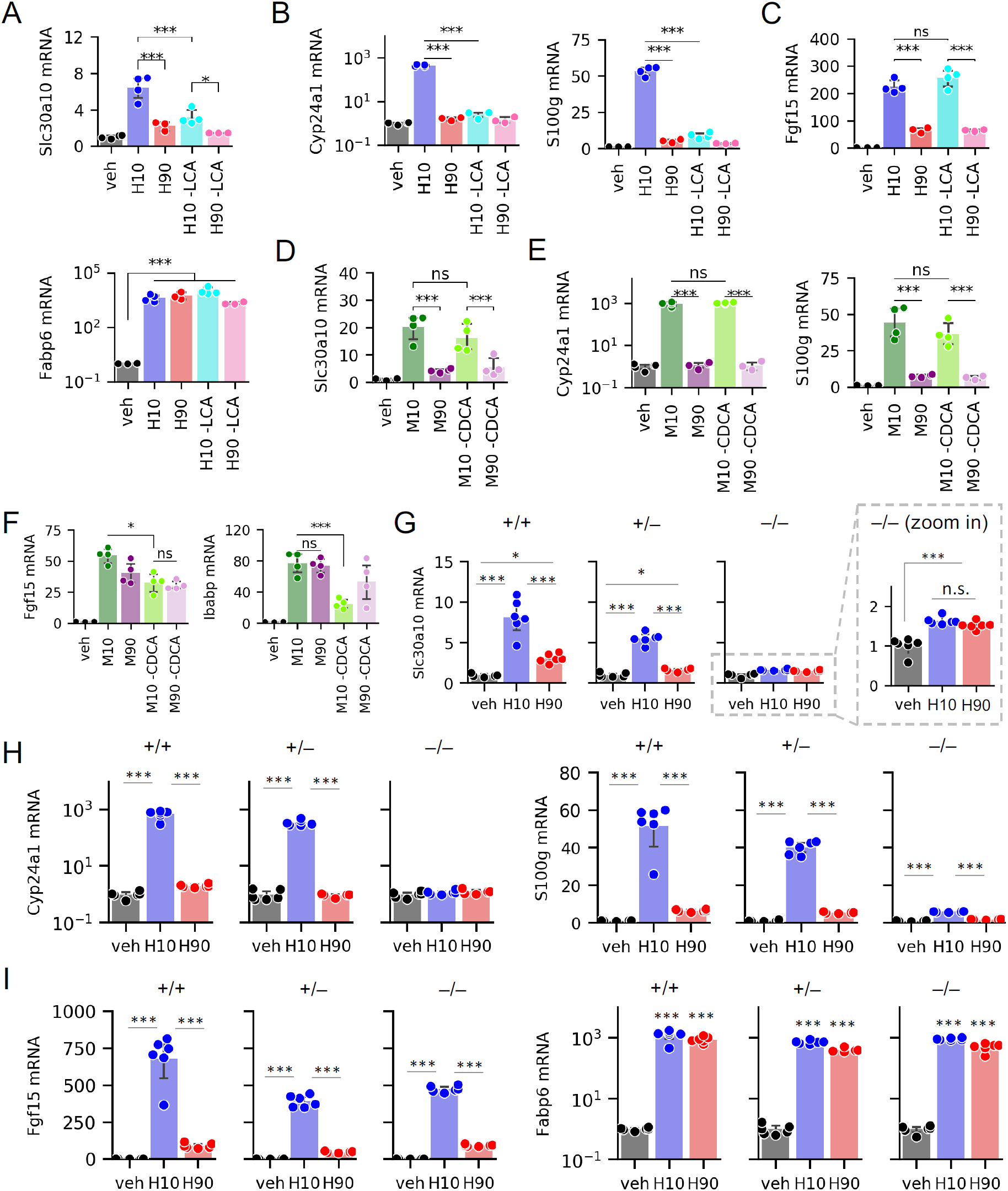
*Slc30a10* expression is regulated by LCA–VDR signaling. (A-E) mRNA levels of *Slc30a10*, VDR targets, and FXR targets in wild-type ileal organoids after treatment with (A-C) human BA pools containing or lacking LCA or (D-F) mouse BA pools containing or lacking CDCA. N=3-4 wells of organoids per condition. (G-I) Expression of *Slc30a10*, VDR targets, and FXR targets in organoids derived from *Vdr*^*+/+*^, *Vdr*^*+/–*^, and *Vdr*^*–/–*^ littermates after exposure to vehicle or human BA pools. n=6 wells of organoids per condition. \**p*<0.05 and \*\*\**p*<0.001.

To test whether CDCA contributes to differential *Slc30a10* induction between low and high 12HBA pools, we treated gut organoids with M10 and M90 pools containing or lacking T-CDCA (Supp. Table 2). In pools lacking T-CDCA, we compensated its removal by increasing other non-12HBAs such that the total BA pool concentrations matched. Removing T-CDCA had no significant effect on *Slc30a10* mRNA levels (Fig. 3D), suggesting CDCA is dispensable for *Slc30a10* induction. Similarly, there was no effect of removing T-CDCA on *Cyp24a1* or *S100g* expression (Fig. 3E). CDCA and its conjugates are potent agonists of FXR (5, 6, 9). Consistent with this, removing T-CDCA from M10 and M90 pools resulted in blunted *Fgf15* and *Fabp6* induction (Fig. 3F). Together, these findings indicate that LCA, but not CDCA, is the BA primarily responsible for the differential expression of *Slc30a10* between low-versus high-12HBA pools.

To directly test the requirement of VDR for BA-induced *Slc30a10* transcription, we carried out experiments on gut organoids derived from *VDR*^*+/+*^, *VDR*^*+/*–^, and *VDR*^−/−^ littermate mice. Strikingly, *VDR*^−/−^ gut organoids showed substantially blunted *Slc30a10* transcription in response to H10 pool (Fig. 3G). Furthermore, *VDR*^+/–^ gut organoids increased *Slc30a10* transcription to intermediate levels between that in *VDR*^*+/+*^ and *VDR*^−/−^ organoids, suggesting a gene dosage effect. As expected, ablating *VDR* diminished the induction of *Cyp24a1* and *S100g* (Fig. 3H), but had little effect on *Fgf15* and *Fabp6* (Fig. 3I). We noted that *VDR*^−/−^ gut organoids still showed a small but significant increase in *Slc30a10* upon H10 and H90 treatment, suggesting possible residual activation by another transcription factor. Thus, we conclude that the induction of *Slc30a10* by low 12HBA pools is largely mediated by VDR.

### Effects of BA composition on other Mn transporters

In addition to SLC30A10, enterocytes express at least two other Mn transporters. The Divalent Metal Transporter 1 (DMT1, encoded by *SLC11A2*) transports Mn from the intestinal lumen into enterocytes (31). SLC39A14 takes up Mn from the basolateral side of enterocytes (50), and it has been reported that SLC39A14 acts synergistically with SLC30A10 to mediate whole-body Mn excretion (50–52). However we found that BA treatments had no effects on *Slc11a2* or *Slc39a14* expression in murine organoids (Fig. 4A), and no effects on *SLC11A2* or *SLC39A14* in Caco-2 cells (Fig. 4B).

**Figure 4.**
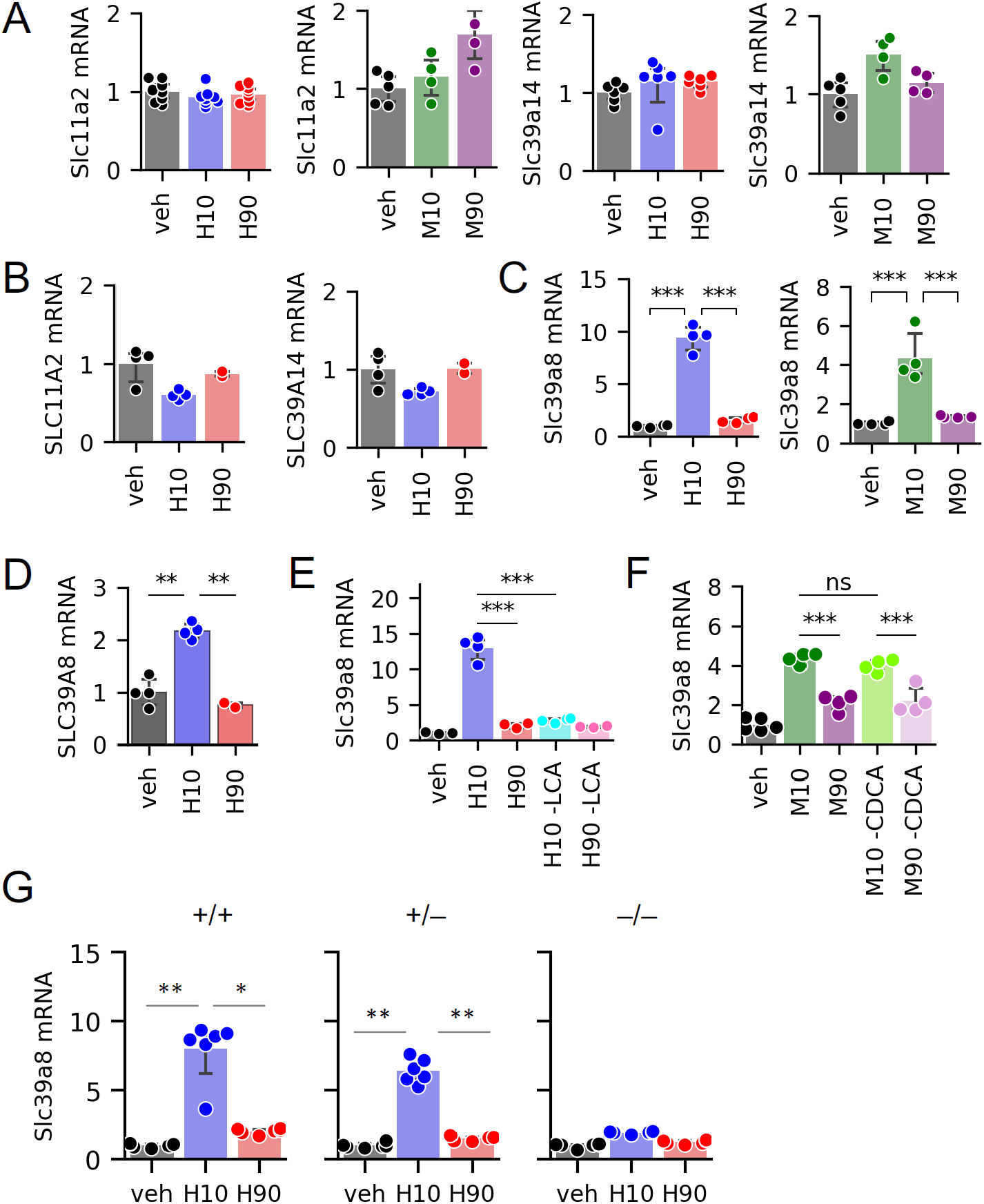
BA effects on additional Mn transporters. Expression of (A) *Slc11a2* and *Slc39a14*, (C) *Slc39a8* in organoids, and (B, D) their corresponding human genes in Caco-2 cells (n=4-8 wells of cells or organoids per group). (E-F) *Slc39a8* expression in organoids exposed to (E) human BA pools with or without LCA and (F) mouse BA pools with or without CDCA (n=4 wells of organoids per group). (G) *Slc39a8* expression in organoids generated from *Vdr*^*+/+*^, *Vdr*^*+/–*^, and *Vdr*^*–/–*^ littermates after exposure to human BA pools (n=6 wells of organoids per group). \**p*<0.05, \*\**p*<0.01, and \*\*\**p*<0.001.

Another Mn transporter is SLC39A8. This protein is highly expressed in liver, where it is critical for Mn uptake from bile into hepatocytes (53). The localization and physiological relevance of SLC39A8 in intestine has not been reported. We found that gut organoids treated with H10 and M10 pools upregulated *Slc39a8* transcription more strongly than H90 and M90 pools (Fig. 4C). Caco-2 cells also showed increased *SLC39A8* expression upon H10, but not H90, treatment (Fig. 4D). Further analyses showed that, similar to *Slc30a10, Slc39a8* transcription was blunted upon removal of LCA from BA pools, and in gut organoids from *VDR*^*–/–*^ mice, whereas removal of CDCA from BA pools had no effect (Fig. 4E-G). These data support the possibility that both Slc30a10 and Slc39a8 are regulated by LCA-to-VDR signaling.

### LCA and vitamin D induce Slc30a10 expression in mouse ileum

To determine the distribution of *Slc30a10* and *Slc39a8* expression *in vivo*, we quantified mRNA expression in murine enterohepatic tissues. We found that all segments of the small intestine expressed *Slc30a10* mRNA at levels higher than the liver (Fig. 5A). We also detected *Slc39a8* expression in small intestine, at levels approximately 10-30% of those in liver (Fig. 5B).

**Figure 5.**
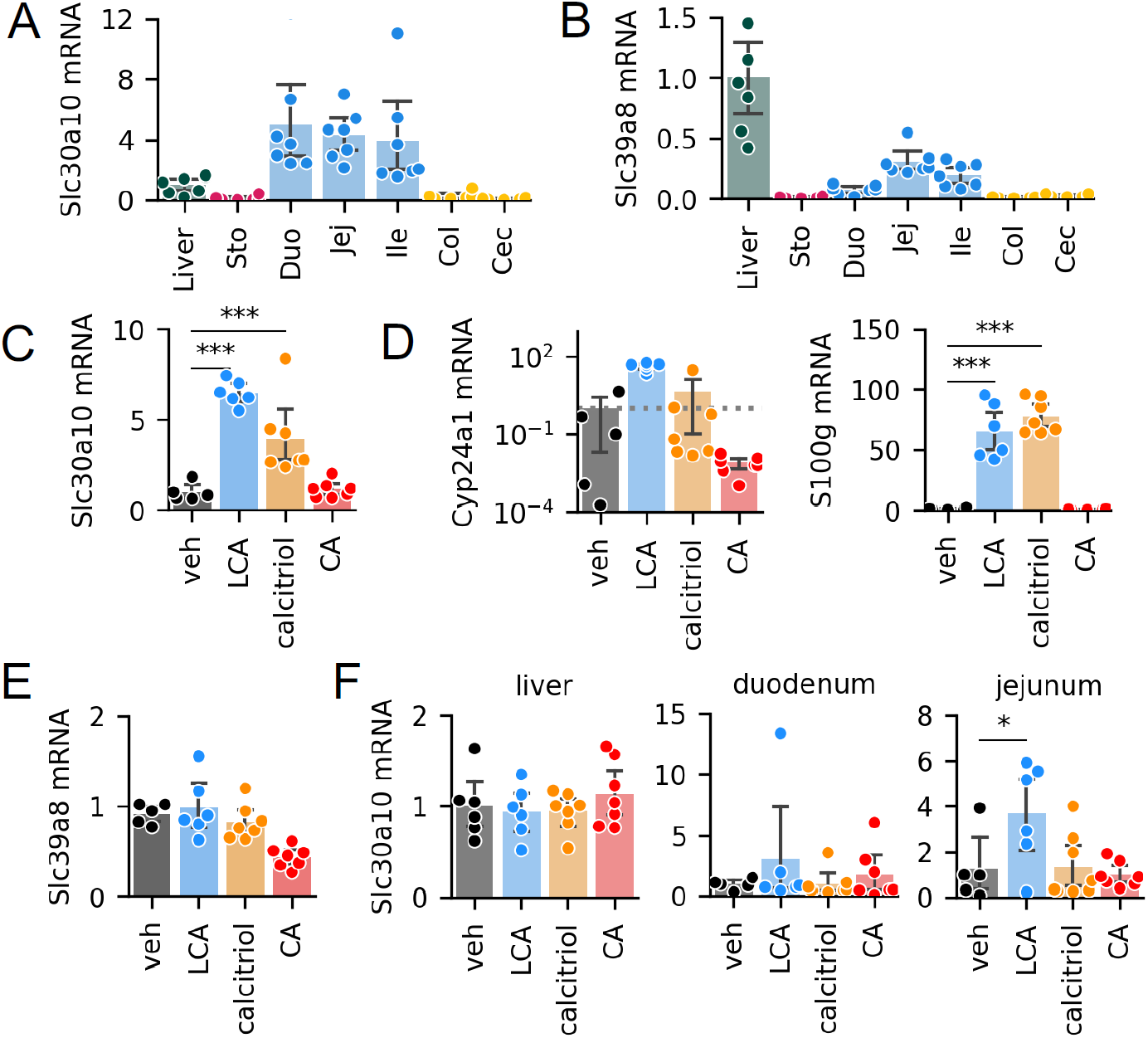
VDR activation increases *Slc30a10* transcription *in vivo*. Expression of (A) *Slc30a10* and (B) *Slc39a8* in liver and gastrointestinal tissues in C57BL/6 mice (n=7). Liver was used as reference tissue. Ileal expression of (C) *Slc30a10*, (D) VDR targets *Cyp24a1* and *S100g*, and (E) *Slc39a8* after oral gavage with corn oil supplemented with vehicle, LCA, calcitriol, or CA. n=5–7 mice per group. (F) *Slc30a10* expression in liver, duodenum, and jejunum after gavage experiment. \**p*<0.05, \*\**p*<0.01, and \*\*\**p*<0.001.

Next, we tested whether LCA and VDR signaling induces expression of Mn transporters *in vivo*. We administered (i) vehicle, (ii) LCA, (iii) the VDR agonist calcitriol, or (iv) the 12HBA cholic acid (CA) to C57BL/6 mice by oral gavage and analyzed mRNA expression in ileal epithelia. Mice receiving LCA and calcitriol showed on average 4- to 6-fold increases in *Slc30a10* expression in ileum (Fig. 5C). The increase in *Slc30a10* expression was accompanied by increased *Cyp24a1* and *S100g* (Fig. 5D), consistent with activation of VDR. Notably, CA did not induce *Slc30a10, Cyp24a1*, or *S100g* (Fig. 5C-D). In contrast to the results of *ex vivo* and *in vitro* experiments, *Slc39a8* was not induced by any of the treatments (Fig. 5E). Interestingly, *Slc30a10* was also induced by LCA in jejunum, but not in duodenum or liver (Fig. 5F). These data show that *in vivo*, LCA promotes *Slc30a10* expression in ileum and jejunum.

## Discussion

Our analyses reveal that BA composition regulates the expression of *Slc30a10* and cellular Mn efflux. SLC30A10 is one of three transporters identified thus far to be critical for whole-body Mn homeostasis in humans and mice (34, 35, 39, 40, 52, 54–56). We found that BA pools low in 12HBAs, which have increased abundance of the non-12HBA LCA, act primarily via VDR to promote *Slc30a10* expression. We also found that ileal *Slc30a10* was inducible by LCA and the VDR agonist calcitriol, indicating that LCA and VDR signaling modulate intestinal Mn transport.

Similar to BAs, a significant portion of Mn undergoes enterohepatic cycling (57, 58). Thus, enterocytes are exposed to luminal Mn from two sources: bile and diet. In adult humans, only about 3-5% of ingested Mn is absorbed (31), indicating robust regulation of intestinal Mn absorption. Obstructing the bile duct results in reduced excretion rates of intravenously administered ^54^Mn in rats (59), highlighting the importance of Mn clearance by the liver and final excretion in the feces. But the intestine appears to play an important role in Mn homeostasis as well, especially when liver excretion has been saturated (60). While deletion of Slc30a10 only in hepatocytes is insufficient to cause hypermanganesemia, combined deletion in liver and gastrointestinal tract causes severe hypermanganesemia and neurotoxicity (39, 41), mimicking the whole-body Slc30a10 knockout mice, though less severe (39, 40). These findings support the notion that intestine and hepatobiliary excretion systems both participate in Mn homeostasis.

Mn is a cofactor for numerous metalloproteins involved in many cellular processes (31, 61). Yet, Mn in excess amounts is toxic. Hypermanganesemia results in dystonia, parkinsonism, polycythemia, and cirrhosis (31, 34, 35, 42, 52, 62, 63). Most cases of Mn toxicity are due to occupational and environmental exposures (31, 62). Mn toxicity has also been reported in cases of severe liver diseases, in patients receiving total parenteral nutrition, and in rare genetic loss-of-function mutations (34, 35, 44, 64–67). Current treatment for Mn toxicity is with chelation therapy, which increases urinary Mn excretion (44), but is nonspecific. Since the identification of disease-causing mutations of *SLC30A10*, there has been considerable progress in understanding how the transporter works and where it is localized in cells (36, 68). However, the regulation of the transporter is incompletely understood.

We identified BA composition, especially LCA abundance, as a modulator of *Slc30a10* expression *in vitro, ex vivo*, and in mouse ileum. We also showed that VDR is the primary mediator of this signaling pathway. This is consistent with a previous report showing VDR activation inducing *SLC30A10* in Caco-2 cells, and that this requires the VDRE in SLC30A10’s promoter region (45). Claro da Silva and colleagues previously reported increased *SLC30A10* mRNA levels in half of duodenal biopsies from volunteers administered oral calcitriol (0.5 µg for 10 days) (45). However, ileal biopsies were not reported, and ileum is expected to be the primary site of uptake of bile acids into enterocytes (49). Our data show that *Slc30a10* expression is induced by LCA and calcitriol *in vivo*, in mouse ileum. Published data from *in vitro* studies using Caco-2 cells and the neuroblastoma cell line SH-SY5Y have also proposed that the ER stress response factor ATF4 and the zinc-sensitive transcription factor ZNF658 contribute to *SLC30A10* regulation (69–71). The identification of this new BA-sensitive pathway may shed light on a known phenomenon: Mn poisoning has been reported in patients on total parenteral nutrition (65–67), suggesting that the lack of intestinal BA signaling leads to reduced SLC30A10 and impaired efflux of Mn from enterocytes into the intestinal lumen, consequently reducing Mn excretion.

In our experiments, *Slc30a10* induction was not completely abrogated upon removal of LCA from BA pools or deletion of VDR. Thus, other BAs and nuclear receptors might also contribute to BA-dependent induction of *Slc30a10*. A published ChIP-seq dataset identified two potential FXR binding sites within 1 kb of the first exon of mouse *Slc3010*, and another potential site within 5 kb (72). Although these binding peaks are approximately an order of magnitude smaller than those seen for classical intestinal FXR targets *Fgf15* and *Slc51a* (72), it is possible that FXR participates in *Slc30a10* regulation at low levels or under certain conditions.

Does BA-dependent regulation of Mn homeostasis have any impact on cardiometabolic disease? Some researchers have reported that patients with type-2 diabetes have plasma or serum Mn at levels higher than controls, while others have reported the opposite (73–76). Because Mn levels must be maintained within a small range (31, 44), it makes sense that both deficiency and excess of Mn would lead to negative consequences. Accordingly, a recent report showed a U-shaped relationship between plasma Mn and the odds ratio for type-2 diabetes (77). When interpreting the previous findings, it should be noted that the most accurate circulating Mn levels are obtained from whole blood, not serum or plasma, since over 60% of blood Mn is found in erythrocytes (78). Interestingly, intravenous glucose tolerance tests in individuals with chronic manganism revealed reactionary hypoglycemia, which was proposed to be due to dysregulation of the hypothalamus-pituitary-adrenal axis (79). Human genetic variants in *SLC39A8* are linked to blood Mn levels as well as body mass index, total cholesterol, and HDL-cholesterol (80–83). In an unusual case of diabetes, a patient showed hypoglycemia in response to oral MnCl_2_, which was abolished upon a partial pancreatectomy (84). Thus although severe Mn deficiency and toxicity lead to overt neurological and liver disorders, the role of Mn in common cardiometabolic diseases might be more nuanced.

Mechanistically, how could Mn contribute to features of cardiometabolic disease? Mn is a required or preferred cofactor for many enzymes, including arginase, phosphoenolpyruvate carboxykinase (PEPCK), and pyruvate carboxylase (PC) (31, 85). Arginase competes with endothelial nitric oxide synthase (eNOS) for their substrate, l-arginine (86). Increased arginase activity has been implicated in endothelial dysfunction in diabetic and atherosclerotic settings, and arginase inhibition has been proposed to improve microvascular complications such as diabetic nephropathy and retinopathy (87, 88). Importantly, arginase activity can be fine-tuned by Mn levels. This was demonstrated in liver extracts from mice rendered Mn-deficient by knockout of the Mn uptake transporter Slc39a8. In these livers, arginase activity was significantly lower than in liver extracts from control *Slc39a8*^*flox/flox*^ mice (53). Conversely, adding MnCl_2_ to the reaction buffer or overexpressing SLC39A8 led to increased arginase activity (53). Interestingly, mice and rats rendered diabetic by streptozotocin injections showed increased liver arginase activity and increased hepatic Mn content (89), suggesting insulin signaling might regulate liver Mn levels and Mn-dependent enzymes. The liver enzymes PEPCK and PC catalyze critical steps for gluconeogenesis. Additionally, oxaloacetate production by PC can also enter other pathways, one of which is the TCA cycle, and reduced PC activity leads to build up of lactate (90, 91). Although Mn deficiency has been reported to reduce activities of these enzymes in adult rats (92, 93), the effects of excess Mn on these enzymes have not been reported. Whether (a) BA regulation of SLC30A10 affects the activities of these Mn-dependent enzymes *in vivo* and (b) these Mn-sensitive enzymes partially mediate the relationship between BA composition and metabolic health are of interest for future investigations.

It has been well-established that BAs, and BA pool composition, regulate macronutrient metabolism. This work demonstrates that BAs and BA pool composition also regulate metal homeostasis, which may be relevant for health and disease.

## Experimental Procedures

### Reagents

Taurine-conjugated α- and β-MCAs were purchased from Cayman Chemical. All other BAs, micelle components, corn oil, and MnCl_2_ were obtained from Sigma. Calcitriol was obtained from Selleck Chemicals. Unless noted otherwise, all cell culture media and supplements were purchased from Gibco.

### Crypt isolation and gut organoid culture

Mouse ileal crypts were isolated following methods developed by the Clevers Lab (30), with minor modifications. We euthanized adult (6-12 weeks old) mice using CO_2_ inhalation followed by cervical dislocation. With the exception of the growth medium, all solutions and Matrigel were kept ice-cold during handling of crypts and organoids. We divided the small intestine into 3 parts of equal length and collected the last segment, adjacent to the cecum. We cut open the ileum longitudinally and rinsed it in PBS. We minced the tissue and washed the pieces by vigorously pipetting in PBS, allowing the pieces to settle, and removing the cloudy supernatant. We repeated this wash step until the supernatant was clear, 4-6 times. We then transferred the pieces into PBS containing 2 mM EDTA and incubated the mixture at 4°C for 30-60 min with gentle shaking. Next, we removed the EDTA-PBS and replaced it with PBS containing 10% fetal bovine serum (FBS). We harvested crypts by vigorously pipetting, allowing the segments to settle, and collected the supernatant in a new tube. We repeated this harvesting procedure with additional FBS-PBS mixture, 3-4 times, pooling the supernatant from the same mouse. We then pelleted the crypts by centrifuging the crypts-FBS-PBS mixture at 900 rpm, 4°C for 5 min. We discarded the supernatant, resuspended the crypts in base medium (Advanced DMEM/F-12 supplemented with 2 mM GlutaMax, 100 U/ml penicillin-streptomycin, and 10 mM HEPES), and pelleted the crypts by centrifuging at 720 rpm, 4°C, 5 min. We discarded the supernatant, resuspended the crypts in base medium, passed the crypts-medium mixture through a 70-µm cell strainer, and centrifuged the crypts-medium mixture at 900 rpm, 4°C, 5 min. Finally, we resuspend the crypts in Matrigel and seeded them on pre-warmed 24-well Nunclon Delta surface-treated plates. After 15-20 min incubation at 37°C, we added IntestiCult Mouse Organoid Growth Medium (STEMCELL Technologies) to the wells. Medium was replaced twice weekly. Mature organoids were passaged every 7-10 days by dissolving the Matrigel dome with vigorous pipetting in PBS, passing the organoids-Matrigel-PBS mixture through a 27 ½- gauge needle, centrifuging to pellet the organoid pieces, resuspending them in new Matrigel, and distributing the organoid-Matrigel mixture into new pre-warmed multiwell plates.

### Caco-2 cell culture

Cells were cultured in DMEM containing GlutaMAX and high glucose, supplemented with 10% FBS and 100 U/ml penicillin-streptomycin and incubated in a 37°C, 5% CO_2_ chamber. Medium was replenished 3 times per week and cells were passaged by trypsinization every 5-7 days.

### Preparation of micelles

We prepared micelles as previously described (94) with minor modifications. Individual stock solutions of oleic acid (OA), 2-palmitoyl glycerol (2-PG), phosphatidylcholine (PC), cholesterol, GUDCA, and LCA were prepared in chloroform. Individual stock solutions of all other BAs were prepared in PBS. We mixed lipophilic components (those in chloroform stocks) in a glass vial and allowed them to dry under N2 stream. We mixed BAs to prepare four distinct BA pools: human low (10%) 12HBA (H10), human high (90%) 12HBA (H90), mouse low (10%) 12HBA (M10), and mouse high (90%) 12HBA (M90). After the lipophilic components dried, we added BA pools to the vials and vortexed the mixture. We further diluted the mixture with DMEM containing 0.2% BSA (base medium), yielding final concentrations of 0.6 mM OA, 0.2 mM 2-PG, 0.2 mM PC, 0.05 mM cholesterol, and 1 mM total BAs. Composition of BA pools are detailed in Table 1 and illustrated in Fig. 1B.

### BA treatments of organoids and cells

We removed gut organoids from Matrigel and exposed the lumens by pipetting in ice-cold PBS followed by centrifugation at 150*g*, 4°C for 10 min. Following a second PBS wash, we placed exposed gut organoids to micelle components without BAs (vehicle), or mixed micelles containing BAs (H10, H90, M10, or M90) in base medium (DMEM, 0.2% BSA) for 24 hours. For BA treatments in the absence of micelles, BA pools were prepared in PBS and diluted in base medium for a final concentration of 1 mM, to match the concentration in the mixed micelle treatments. For Caco-2 cells, we seeded 200,000 cells/well into 12-well plates and allowed cells to reach confluency. We replaced the growth medium, washed cells with PBS, added test media to the cells as indicated in the experiments.

### RNA sequencing

We extracted RNA using the RNeasy Mini Kit (Qiagen). RNA concentration and quality were determined by Qubit Bioanalyzer. RNA from 3 samples per treatment group were submitted to the JP Sulzberger Columbia Genome Center for library preparation (standard poly-A pull-down for mRNA enrichment), RNA sequencing (30M depth, single read), and processing of raw data. Differential expression analysis on counts data was done using the DESeq2 package in R.

### Mn efflux assays

Mn efflux assays were done as previously described (36). We collected organoids by dissolving the Matrigel dome with ice-cold PBS, pipetting, and centrifuging at 150 *g* for 10 min. Organoids were first incubated in ‘exposure media’ (DMEM supplemented with 0.2% BSA containing 0 or 500 µM MnCl_2_) for 16h at 37°C, 5% CO_2_. Following exposure, organoids were washed with PBS, given media containing micelle components (vehicle), 1 mM H10, or 1 mM H90, and incubated for another 20h. Organoids were pelleted by centrifugation and efflux media were collected in metal-free tubes. For Mn efflux assay in Caco-2 cells, we followed the same procedure, with the exception of 100 µM MnCl_2_ for exposure. Mn was measured by inductively coupled plasma mass spectrometry (ICP-MS).

### Mouse experiments

12 week-old male C57BL/6 mice (Taconic) were given AIN-93G with vitamin D adjusted at 50 IU/kg for 3 days before the experiment. Mice were housed in a facility with 12h light/12h dark cycle and had free access to food at all times. Ethanol (10%, vehicle), 0.8 mmol/kg LCA, 50 nmol/kg calcitriol, and 0.8 mmol/kg CA were dissolved in corn oil. Oral gavages were done at 14 h and 2 h prior to euthanasia. Experiments were approved by the Institutional Animal Care and Use Committee of Columbia University Medical Center. Doses and time points were based on previous studies by Ishizawa and colleagues (95). We collected ileum epithelia by cutting open the segment longitudinally and scraping the mucosa layer away from the muscle wall. Tissues were stored at –80°C until processed for RNA extraction.

### RNA extraction, cDNA synthesis, and qPCR

We extracted RNA from gut organoids using the RNeasy Mini Kit (QIAGEN) and used 200 ng for cDNA synthesis. We extracted RNA from Caco-2 cells and ileum epithelium scrapings using TRIzol (Thermo Fisher) and used 1 µg for cDNA synthesis. cDNA synthesis was done using the High Capacity cDNA Reverse Transcription Kit (Thermo Fisher). Quantitative real-time PCRs were carried out using iTaq Universal SYBR Green Supermix (Bio-Rad) and the CFX96 Real-Time PCR detection system (Bio-Rad). Mouse *36b4* or *B2m* were used as for reference genes for samples from mouse gut organoids and tissues. Human *RPLP0* (primers from QIAGEN) was used as reference gene for cDNA from Caco-2 samples. Primer sequences are listed in Supp. Table 3.

### Statistical analysis

Unless noted otherwise, we used one-way ANOVA followed by Benjamini-Hochberg correction. Adjusted p-values less than 0.05 were considered significant, with \**p*<0.05, \*\**p*<0.01, and \*\*\**p*<0.001 between groups as marked.

## Acknowledgements

The authors would like to acknowledge Ana Flete and Thomas Kolar for technical assistance. This work was supported by funding from the NIH–DK115825 to RAH and T32DK007328 to TRA–and the Russell Berrie Foundation. This research was funded in part through the NIH/NCI Cancer Center Support Grant P30CA013696 and used the Genomics and High Throughput Screening Shared Resource.

## FOOTNOTES

The content is solely the responsibility of the authors and does not necessarily represent the official views of the National Institutes of Health

## Abbreviations

12-HBA: 12α-hydroxylated bile acid;
α-MCA: α-muricholic acid;
β-MCA: β-muricholic acid;
BA(s): bile acid(s);
CA: cholic acid;
CDCA: chenodeoxycholic acid;
DCA: deoxycholic acid;
FXR: farnesoid X receptor;
LCA: lithocholic acid;
UDCA: ursodeoxycholic acid;
VDR: vitamin D receptor

