## Supporting Information for "Bile acid composition regulates the manganese transporter Slc30a10 in intestine"

This PDF file includes:

Tables S1 to S3
Supp. Fig. 1

**Table S1. Composition of BA pools with and without LCA**

| BA group | BA species | % in H10 | % in H90 | % in H10 –LCA | % in H90 –LCA |
|---|---|---|---|---|---|
| 12HBA | G-CA | 7 | 63 | 7 | 63 |
| 12HBA | G-DCA | 3 | 27 | 3 | 27 |
| Non-12HBA | G-CDCA | 86.85 | 9.65 | 87.7 | 9.75 |
| Non-12HBA | G-UDCA | 2.25 | 0.25 | 2.3 | 0.25 |
| Non-12HBA | LCA | 0.90 | 0.10 | 0 | 0 |

G- = Glycine-conjugated

**Table S2. Composition of BA pools with and without CDCA**

| BA group | BA species | % in M10 | % in M90 | % in M10 –CDCA | % in M90 –CDCA |
|---|---|---|---|---|---|
| 12HBA | T-CA | 9.0 | 81.0 | 9.0 | 81.0 |
| 12HBA | T-DCA | 1.0 | 9.0 | 1.0 | 9.0 |
| Non-12HBA | T-CDCA | 13.5 | 1.5 | 0 | 0 |
| Non-12HBA | T-UDCA | 8.1 | 0.9 | 8.1 | 0.9 |
| Non-12HBA | LCA | 0.9 | 0.1 | 0.9 | 0.1 |
| Non-12HBA | T-α-MCA | 22.5 | 2.5 | 27.0 | 3.0 |
| Non-12HBA | T-β-MCA | 45.0 | 5.0 | 54.0 | 6.0 |

T- = Taurine-conjugated

**Table S3. Primer sequences**

| Gene | Primer 1 | Primer 2 |
|---|---|---|
| **Mouse primers** | | |
| *36b4* | AGATGCAGCAGATCCGCAT | GTTCTTGCCCATCAGCACC |
| *Cyp24a1* | CCAAGGTCCGTGACATCCAA | GATGCACCGAGTCGAAGGAG |
| *Fabp6* | GACGTGATTGAAAGGGGACG | CTCATCTTCACGGTAGCCT |
| *Fgf15* | ACGGGCTGATTCGCTACTC | TGTAGCCTAAACAGTCCATTTCCT |
| *S100g* | GCTGTTCCTGTCTGACTCCT | GCTGGGGAACTCTGACTGAA |
| *Slc11a2* | CAGGAAGTCATTGGCTCAGC | TATCCAAACGTGAGGGCCAT |
| *Slc30a10* | GAGATGGGCCGTTACTCAGG | GCCTCCACGAAGATGGTGAA |
| *Slc39a14* | ATCCAGAATCTTGGCCTCCT | AAGAGCTGCCTTTTCCATGA |
| *Slc39a8* | CTGTCACTGAGCCTAACGGA | GCCGTCGATGAAATTGTGGA |
| *Slc51b* | GAAGATGCGGCTCCTTGGA | GCTCTCTGTTTCCTGTGGGT |
| **Human primers** | | |
| *RPLP0* | QIAGEN PPH21138F-200 | |
| *CYP3A4* | GCAGGAGGAAATTGATGCAGTT | GTCAAGATACTCCATCTGTAGCACAGT |
| *CYP24A1* | GGGTCTCAAGAAACAGCACG | GCCTTCCACGGTTTGATCTC |
| *FABP6* | ACTTGGTCCCAGCACTACTC | TAGGCCAGTCTCTTGCTCAC |
| *FGF19* | CTGACATGTTCTCTTCGCCC | CGTGGACTCAGGACTGTTCT |
| *SLC11A2* | TGCATCTTGCTGAAGTATGTCACC | CTCCACCATCAGCCACAGGAT |
| *SLC30A10* | TTCAACATGCTCTCCGACCT | GAAGATGGTGAAGCAGAGCG |
| *SLC39A14* | TTCACCCCTGGCATTAGCAG | CACCCGTGGGATTCTCAACA |
| *SLC39A8* | TGCCTGGATGATAACGCTCT | AGCCCAACATAGCAGGAACA |

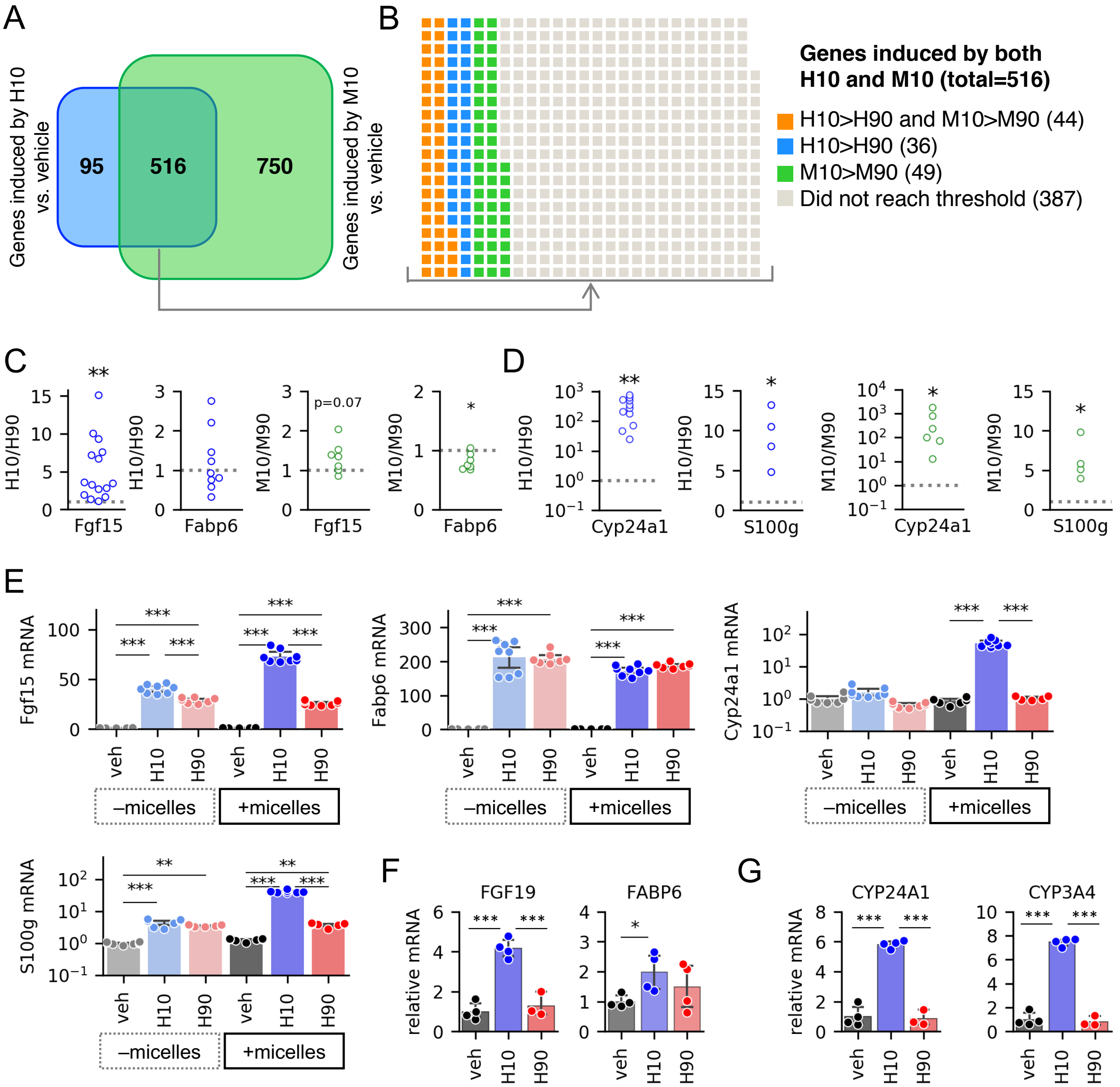


**Supporting Figure 1. Validation of differential expression of FXR and VDR target genes.** (A) Venn diagram of differentially expressed genes induced by H10 and/or M10 compared to vehicle. (B) Genes preferentially induced by low 12HBA pools. Grouped based on $\log_2$(expression in H10/expression in H90)>1 or log2(expression in M10/expression in M90)>1 and $p_{adj}$<0.05. (C, D) Ratio of gene expression induced by low 12HBA to high 12HBA. Each point represents the data derived from a different mouse (n=7-15). The data point plotted is the ratio of (average mRNA levels in low 12HBA-treated organoids [n=3-8 wells])/(average mRNA levels in high 12HBA-treated organoids [n=3-8 wells]). Ratios >1 signify higher expression in low 12HBA-treated group, ratios <1 signify higher expression in high 12HBA-treated group, ratios = 1 signify no difference between low- versus high- 12HBA-treated groups. The gray dotted line marks ratio = 1. *$p$<0.05, **$p$<0.01 one-sample t-test vs. μ=1. (C) FXR targets, (D) VDR targets. (E) Gene expression in gut organoids treated with BA pools in the absence or presence of micelles. N=8 wells of organoids per group. (F, G) Gene expression of FXR and VDR targets in Caco-2 cells. N=3-4 wells of cells per group. **$p$<0.01 and ***$p$<0.001.
